# Deep Learning Based Cross-Modality Histological Brain Section Registration in Multiple Species Using Synthetic Images

**DOI:** 10.64898/2026.07.17.738987

**Authors:** Siqi Fang, Daniel Tward

## Abstract

Large-scale brain mapping increasingly relies on integrating histological imaging datasets within standardized anatomical reference frameworks. To accomplish this, registration techniques are used to align imaging datasets between different modalities or to reference atlases. Deep learning approaches have emerged a fast and scalable framework for approaching this challenge, but such methods typically require large annotated datasets that are unavailable in this setting. To address this, we developed a framework for training a convolutional neural network for this task using entirely simulated data. We show that the same approach can be used for different species (mouse and marmoset), and we provide accuracy validation in terms of Dice and Hasudroff distance between anatomical regions in comparison to an alternative method. This approach has the potential to accelerate large or high throughput studies of brain anatomy.

## 1 Introduction

Large-scale brain mapping increasingly relies on integrating histological imaging datasets within standardised anatomical reference frameworks. Registering two-dimensional histological sections to three-dimensional reference atlases enables consistent localisation of anatomical structures, comparison across experiments, and large-scale quantitative analysis of brain organisation. In mouse, the Allen Mouse Common Coordinate Framework (CCF) has become the dominant reference atlas, providing a high-resolution anatomical template and hierarchical region annotations that support data integration across laboratories [24]. Recent work has further extended atlas resources through improved anatomical coverage and population-averaged templates [1], while similar efforts have produced population-based atlases for other species such as the marmoset [10]. These frameworks provide a critical foundation for modern systems neuroscience by enabling spatially standardised analysis of diverse datasets.

A variety of software ecosystems have also been developed to support atlas alignment workflows. Although these systems provide flexible and powerful workflows, they typically rely on semi-automated interaction and multiple processing steps, which can limit scalability for large datasets. The EBRAINS registration suite provides tools such as QuickNII for interactive affine alignment and VisuAlign for manual or semi-automated deformable refinement[18,19]. These tools have been widely adopted for mapping histological datasets into atlas space and are often integrated into broader analysis pipelines. For example, the ABBA framework incorporates DeepSlice-based initialisation within a Fiji-based workflow that supports atlas alignment, segmentation, and whole-brain quantification of gene expression patterns [7].

Despite these advances in atlas construction, aligning histological sections to reference atlases remains a major computational challenge. Histological datasets often consist of serial two-dimensional sections acquired using different staining protocols, imaging modalities, and experimental conditions. Tissue distortions introduced during slicing, mounting, and imaging further complicate registration. Classical deformable image registration methods such as diffeomorphic registration and symmetric normalisation (SyN) implemented in the ANTs framework have been widely used for anatomical alignment tasks in neuroimaging [3]. While these methods provide highly accurate spatial transformations, they are computationally expensive and typically require iterative optimisation, making them less suitable for large-scale high-throughput histological pipelines.

Therefore, recent work has explored machine learning approaches for automated slice localisation. DeepSlice demonstrated that convolutional neural networks can predict the spatial position of mouse brain sections within a reference atlas by regressing a set of anchor vectors describing the orientation and position of the slice [6]. This approach achieves localisation accuracy comparable to human experts while reducing the time required for registration by several orders of magnitude. Other systems such as AMBIA adopt a modular architecture in which deep learning-based slice localisation is followed by deformable registration using classical methods such as LDDMM [20]. More broadly, deep learning approaches for medical image registration have emerged as a promising alternative to classical optimisation-based methods, with frameworks such as VoxelMorph demonstrating that neural networks can directly predict deformation fields for fast image alignment [4]. One thing to note that is methods such as AMBIA treat localisation and deformable registration as independent steps and may still require manual interaction or landmark placement for optimal performance.

Cross-modality variability further complicates automated registration. Brain images can differ dramatically in contrast and appearance depending on staining protocols, imaging modalities, and acquisition pipelines. For example, Nissl staining, immunohistochemistry, autofluorescence imaging, and serial two-photon tomography all produce images with distinct visual characteristics. Several studies have attempted to address this problem using image translation techniques based on generative adversarial networks, such as CycleGAN or Pix2Pix, which transform images from one modality into another prior to registration [12]. Domain adaptation approaches provide alternative strategies for learning modality-invariant representations, including adversarial training and distribution alignment methods [11].

Another limitation of existing slice registration systems is their restricted scope across species and experimental contexts. Most automated slice-localisation tools have been developed specifically for the mouse brain and rely on assumptions tied to the Allen CCF atlas. Meanwhile, atlas alignment pipelines for other species such as the marmoset often rely on classical registration algorithms such as ANTs and require substantial manual configuration [10]. Developing methods that generalize across species and atlas frameworks would facilitate broader adoption and improve interoperability between experimental datasets.

In this work, we present a fully automated pipeline for registering two-dimensional coronal histological brain sections to three-dimensional reference atlases, as summarized in Fig. 1. First, our approach integrates deep learning-based slice localization with learned affine and deformable registration, enabling end-to-end alignment without manual intervention. Unlike previous approaches (AMBIA) that rely on classical deformable registration algorithms, our method predicts deformation fields directly using convolutional neural networks, enabling fast registration in a single forward pass. Second, to address cross-modality variability, we introduce a training strategy based on progressive colour augmentation combined with a differentiable image similarity loss that operates on atlas-derived reference slices, similar to the approach used in SynthMorph[14]. This approach encourages the network to learn modality-invariant representations without requiring explicit modality translation or paired training data. In addition, our framework supports multiple species by using the same architecture to register both mouse brain sections to the Allen CCFv3 atlas (which contains a population average serial two photon image, and a single subject Nissl image) and marmoset sections to the Brain/MINDS atlas [25] (which contains a single subject MRI and Nissl images). Finally, we provide a comprehensive evaluation of our method using region-level metrics including Dice similarity and Hausdorff distance across multiple anatomical ontology levels. By transforming atlas annotations back into the input image space, we ensure fair comparison across methods operating in different coordinate systems. Together, these contributions provide a scalable and fully automated solution for atlas alignment of histological brain datasets, enabling efficient integration of large-scale neuroanatomical data across imaging modalities and species.

**Fig. 1:**
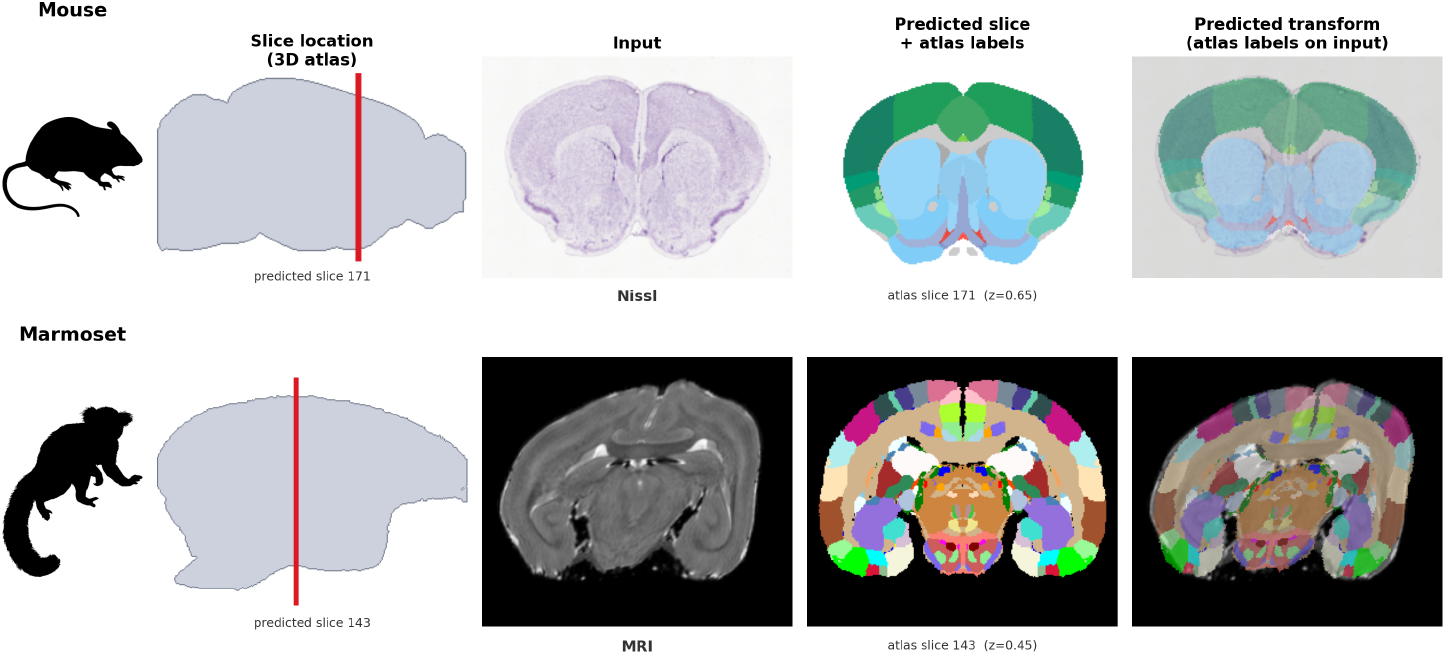
Method overview across species and modalities. Given a single 2D section, our pipeline (i) localises it within a 3D atlas and (ii) predicts a dense transformation that carries atlas region labels onto the section. Top: mouse (Nissl histology); bottom: marmoset (MRI). *Slice location (3D atlas):* the predicted antero-posterior position shown as a plane through a sagittal view of the atlas. *Input:* the 2D section presented to the network. *Predicted slice + atlas labels:* the atlas slice retrieved at the predicted location with its ontology-coloured region labels, in the standard atlas orientation. *Predicted transform:* the estimated transformation warps the atlas labels onto the input, following the section’s own un-normalized anatomy. The method generalises across imaging modalities (Nissl, MRI) and species (mouse, marmoset) with a single architecture.

## 2 Methods

### 2.1 Ontology

For each atlas dataset (mouse and marmoset) we used the hierarchical brain ontology distributed with the atlas. To support multi-scale anatomical analysis, each atlas ontology was parsed into a hierarchical graph. Voxel-wise metrics were computed at each level, where a structure at a coarser level is the union of its descendants at finer levels.

For mouse data, we utilised CCF’s curated st_level field [24]. Conversely, marmoset levels were assigned based on raw tree depth from the root [10]. Because semantic granularity and nesting depth encode different organizational principles, these level numbers are not directly comparable across species. For instance, gross anatomical structures grouped at Mouse level 5 are distributed across Marmoset depths 2 (Midbrain, Pons, Medulla), 3 (Thalamus, Hypothalamus), and 4 (Striatum, Hippocampus, Isocortex). Consequently, results are evaluated within-species at each level rather than between species at identical level numbers. An illustration of the ontologies we used at multiple levels of granularity are shown in Fig. 2.

**Fig. 2:**
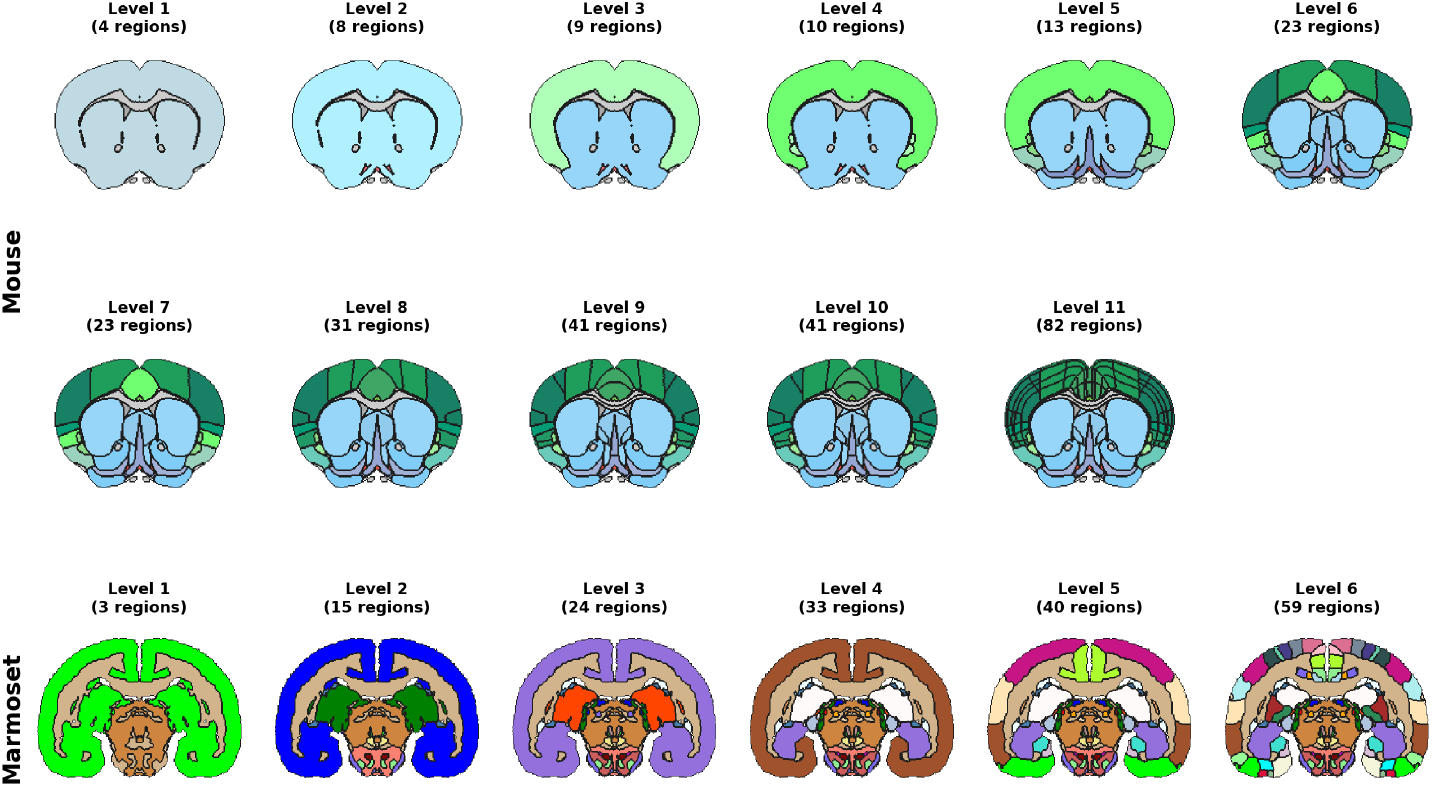
Hierarchical anatomical ontologies used for multi-scale evaluation. A single coronal slice per species (mouse: Allen CCF, slice 171, top two rows; marmoset: Brain/MINDS, slice 143, bottom row) shown at successively finer levels of its region ontology. At each level *L*, every voxel is assigned to its ancestor region at depth *L* in the hierarchy, from a few coarse compartments (cortex, subcortex, fibre tracts) to dozens of fine nuclei and cortical areas. The mouse ontology is substantially deeper (11 levels, up to 82 regions on this slice) than the marmoset one (6 levels, up to 59 regions).

### 2.2 Model training with synthetic data

#### Synthetic Slice Generation

To enable supervised training without manual microscopy labels, 2D slices *S* and label maps are sampled from a 3D atlas. We apply two levels of independent affine and deformable transformations: first warping *S* into *I* and then further warping into *J* (*S → I → J* ). At inference, *I* is the input section and *J* the retrieved atlas slice: the atlas is the *source* of anatomical labels and the input is the registration *target*. The network estimates the transform *T*_*I→J*_ that aligns the input onto the atlas, and its inverse 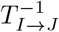 warps the atlas (source) labels into the input (target) space, where all evaluation is performed. Affine warps include random rotations, anisotropic scaling, shear, and translations. Non-linear deformations are generated by smoothing random Gaussian velocity fields, supported on image gradients as described in [17], with a Laplacian regularising kernel [5], then integrating them via scaling-and-squaring [2] to obtain diffeomorphic displacement fields; the same kernel and integration scheme are used by the registration network and detailed in Sec. 2.3. This process yields training pairs (*I, J* ) with exact ground-truth affine matrices *A*_*I→J*_ and velocity fields *v*_*I→J*_ . Applying these random transformations to produce simulated images often places part of the tissue out of the image’s field of view, which may help to build robustness for registering images with some missing tissue.

#### Synthetic RGB Appearance Modelling

(Fig. 3a) To emulate histological modalities like Nissl and Serial Two Photon tomography (STP), Red-Green-Blue (RGB) images are generated from atlas label maps using tissue-specific colouring and noise. Anatomical classes, including grey matter, white matter, and CSF, are assigned either fixed base colors or randomized RGB triplets during augmentation phases. We introduce intensity and texture variability through spatially correlated Gaussian noise, mild blurring to simulate microscope optics, and independent per-pixel noise. Final images are normalized to [0, 1], creating pseudo-microscopy that facilitates modality-invariant registration while preserving anatomical integrity.

**Fig. 3:**
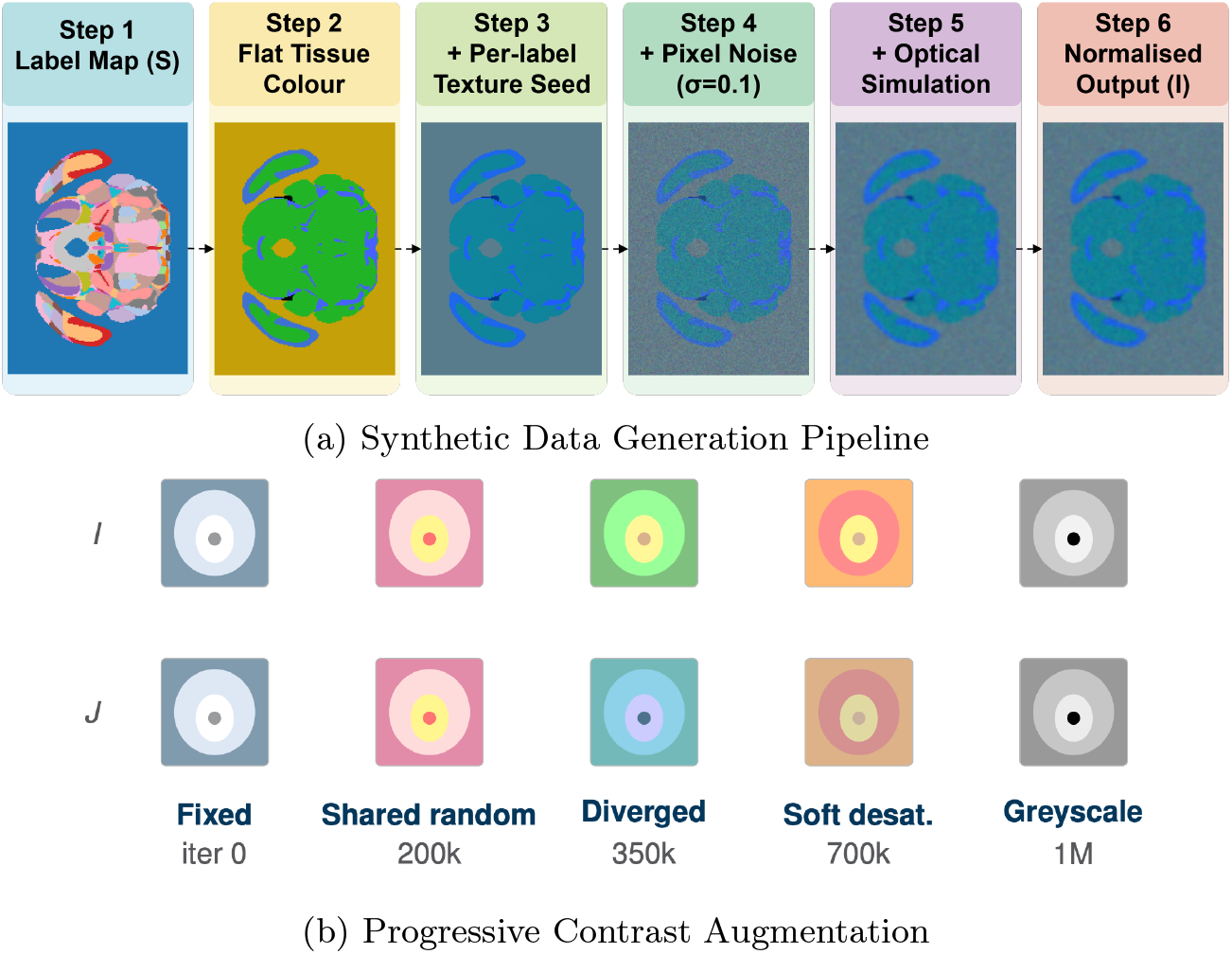
Synthetic Data Generation. (a) Steps to generate a synthetic imge given a set of atlas labels. (b) Our image synthesis curriculum from simple to difficult registration tasks.

#### Progressive Contrast Augmentation

(Fig. 3b) Training the network directly with the full range of contrast randomisation led to poor convergence: with simultaneous geometric and appearance variability, the model failed to establish reliable spatial correspondences early on. We therefore adopt a staged curriculum that increases task difficulty over time, starting from a simple geometry-only setting and progressing towards full modality-invariant alignment. The model first learned geometric alignment using a deterministic colour scheme (first 200*K* image pairs). At 200*K*, colour randomisation was introduced gradually: *I* and *J* within each pair shared the same randomly-sampled colour palette, so contrast between them was preserved while absolute colours varied. At 350*K, I* and *J* were assigned independent random palettes, forcing the network to find spatial correspondences across true cross-modality appearance differences. Beyond 700*K* pairs, training included softer desaturation for 20% of samples. Finally (after 1*M* pairs), a probabilistic ramp (up to 0.5) for hard grayscale conversion accommodated single-channel histological data.

### 2.3 Two-Stage Registration Pipeline

We propose a fully automated end-to-end framework that operates in two stages: (1) Slice Localisation, which identifies the anatomical position of an unknown section, and (2) Deformable Registration, which performs fine-scale deformable alignment to the retrieved atlas slice (Fig. 4).

**Fig. 4:**
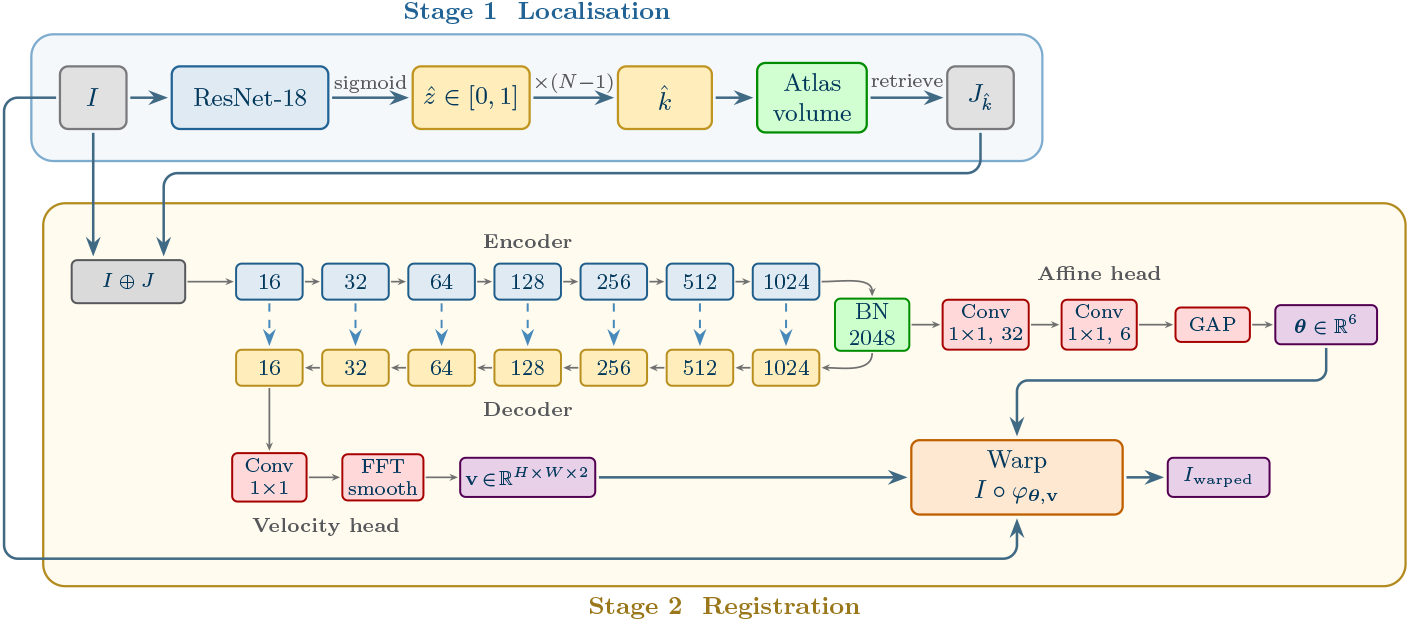
Two-Stage Localisation and Registration Architecture

#### Stage 1: Slice localisation model

Localisation is formulated as a continuous coordinate regression problem to avoid the instability of discrete classification between near-identical adjacent slices. We utilize a ResNet-18 [13] backbone trained from scratch to predict a normalized coordinate 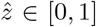. The model is optimized using a Huber (smooth-*L*_1_) loss [15] on the normalized coordinate, with transition point *β* = 0.05: errors smaller than *β* are penalised quadratically (an MSE regime), while larger errors incur only a linear penalty, bounding the gradient for gross mispredictions. As 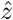 spans [0, 1] over the full anteroposterior range, *β* = 0.05 is 5% of that range, i.e. ≈11 slices for the mouse atlas and ≈16 for the marmoset. The converged localisation errors are far smaller than *β* and thus lie well inside the quadratic regime, so the robust linear regime engages only for the large outlier errors typical of early training. The model is trained in a self-supervised manner on atlas slices transformed with randomised non-linear warps and synthetic appearance variability. We use the same progressive curriculum as the registration model, applied to a single image (no *I, J* pair): training begins with a deterministic colour scheme, progresses to fully randomised colours, and finally introduces soft desaturation and hard grayscale conversion to bridge to single-channel histological data. Because the localiser converges faster, curriculum stage transitions occur at proportionally smaller sample counts (20K, 35K, and 100K image pairs). At inference, the predicted 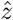 is converted to a discrete atlas index 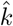 to retrieve the corresponding atlas slice 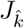 . Because training images are synthesised on the fly with continuously-valued random transformations, the model effectively never encounters the same image twice. We therefore define our *held-out* evaluation set as 256 synthetic samples drawn once from the same generative process with an independent random seed, used solely for evaluation and excluded from gradient updates.

#### Stage 2: Deformable Registration

The registration network utilises a U-Net architecture with seven layers to jointly estimate global and local alignment in a single forward pass. The input section *I* (target) and the retrieved atlas slice 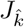 (source) are concatenated as a 6-channel input.

Bottleneck features are globally average-pooled to regress six parameters for a 2 × 3 affine matrix, capturing rotation, scaling, and translation. The deformation decoder predicts a stationary velocity field *v*. To ensure physical plausibility, we enforce diffeomorphic regularity via Laplacian smoothing [5]:

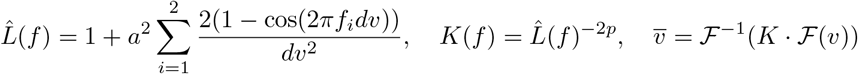

where *f* = (*f*_1_, *f*_2_) is the spatial-frequency vector (cycles *µ*m^−1^) and *F, F*^−1^ denote the two-dimensional discrete Fourier transform and its inverse; *a* = 500 *µ*m sets the smoothing length-scale, *dv* = 50 *µ*m is the voxel spacing, and *p* = 2. With *p >* 1 the smoothing operator 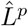 is strong enough that the admissible velocity fields form a reproducing-kernel Hilbert space that embeds continuously into *C*^1^; by the theory of Dupuis et al. [9] the flows they generate are then guaranteed to be diffeomorphisms. The smoothed velocity field *v* is then integrated using the scaling-and-squaring method (*n* = 3) [2] to yield an invertible and differentiable deformation field. This spectral approach provides computational efficiency compared to spatial Gaussian smoothing.

#### Training and Validation

The registration stage is trained on synthetic pairs (*I, J* ) with known transformations, supervised by a weighted sum-of-squares loss on the affine and velocity components: 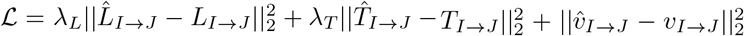, where the affine matrix *A*_*I→J*_ = [ *L*_*I→J*_ | *T*_*I→J*_ ] splits into its 2 × 2 linear part *L* and translation *T* . The weights *λ*_*T*_ = *HW* (the number of pixels) and 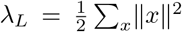 (half the summed squared pixel coordinates) rescale the affine error into the total squared pixel displacement it induces, placing the affine term on the same scale (summed squared displacement over the *H* × *W* grid) and order of magnitude as the per-pixel deformation term 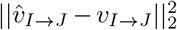, so that neither dominates. This explicit supervision allows the network to learn geometric consistency across large transformations and fine-scale deformations.

Following synthetic training, the unified pipeline is validated on real histological and MRI datasets across mouse and marmoset species, covering nine distinct within- and cross-modality tasks. Four of these tasks are illustrated in Fig. 5.

**Fig. 5:**
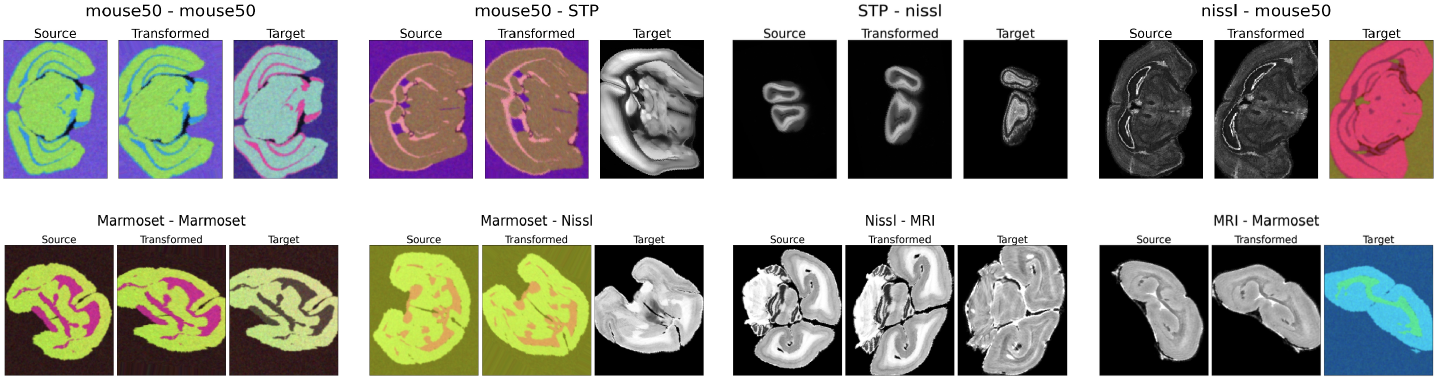
Example validation results showing the input section, the transformed (registered) input, and the atlas slice. Top: Mouse; Bottom: Marmoset.

### 2.4 Evaluation Protocol

The network predicts the forward transform *T*_*I→J*_ aligning the input section (the target) to the atlas (the source of labels). To evaluate error in the native target space, and for fair comparison with DeepSlice which predicts this inverse mapping directly, the atlas (source) labels are carried into the input space via the inverse: 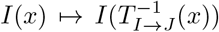. Registration accuracy is evaluated using volume-weighted Dice coefficients and boundary-weighted Hausdorff distances. For a structure *r* at a given ontology level, volumetric overlap is the Dice coefficient 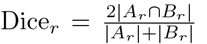, where *A* is the target label map (the input section’s ground-truth labels) and *B* the warped source (the atlas labels registered into the target space), and *A*_*r*_, *B*_*r*_ are the pixel sets of structure *r* in *A* and *B* respectively. Boundary agreement is summarised, per structure, by the 50th and 95th percentiles (*H*_0.95_) of the symmetric distances between the *A*_*r*_ and *B*_*r*_ boundaries.

We compute these per-structure metrics for every structure, at every ontology level, on every slice, and report a single number per level. For overlap this is the *Micro Dice*: a weighted average of the per-structure Dice over all structures at that level on each slice, weighting each structure by its average region size (the mean of its *A*_*r*_ and *B*_*r*_ pixel counts). Equivalently, the intersections and region sizes are pooled across those structures before forming the ratio. We use the term “micro” here, which was introduced in the python package scikit learn as one of several options for aggregating data to produce a single measure. The boundary metric is summarised analogously, as a weighted mean of the per-structure percentiles weighted by each structure’s total boundary length, not a single percentile recomputed from pooled boundary distances. These summaries are presented at each level across the hierarchy of the Allen and Brain/MINDS ontologies, characterising performance from fine-grained nuclei to coarse brain divisions.

To isolate the impact of slice localisation errors, we compare two scenarios: (i) Oracle Selection: Registration using the ground-truth atlas index *k*_*true*_. (ii) Predicted Selection: Registration using the 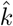 predicted by the Stage 1 network. This protocol quantifies how localisation inaccuracies propagate into the final geometric alignment.

## 3 Results

### 3.1 Registration

For mouse, we evaluated accuracy using three modalities of data from the Allen CCF: Nissl, STP, and randomly coloured atlas labels. Importantly, the first two have not been seen during training, and for the third the same instance of random colors is never seen twice. For marmoset, we evaluated accuracy using three modalities of data from the Brain/MINDS atlas: Nissl, MRI, and randomly coloured atlas labels. Again, the first two have not been seen during training, and for the third the same instance of random colors is never seen twice.

For the mouse dataset, within-modality registration consistently outperforms cross-modality across all ontology levels (Fig. 6). At the gross anatomy level (level 5), the gap is modest but widens at finer scales. This indicates that cross-modality alignment disproportionately affects small and spatially compact structures. Registration performance is approximately symmetric with respect to modality direction, although small asymmetries are observed. In particular, Nissl images are consistently more challenging as targets, suggesting that appearance mismatch plays a stronger role than transformation direction. A non-monotonic dip in performance occurs at level 7 across all modality pairs. This level contains heterogeneous and spatially compact structures (e.g., thalamic nuclei, brainstem regions), which are inherently difficult to align. Importantly, this behaviour is consistent across all methods and conditions, indicating that it reflects atlas structure rather than model failure.

**Fig. 6:**
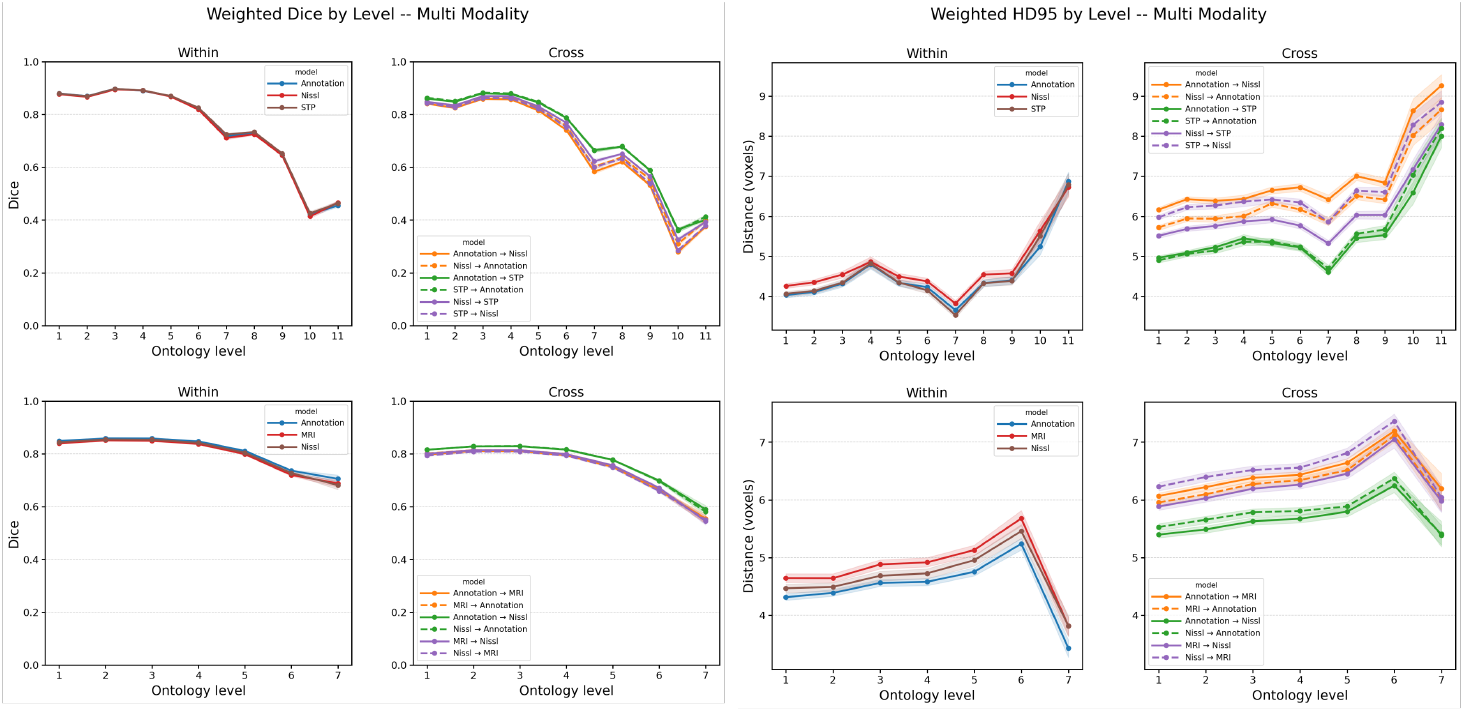
HD95 (right) and Micro Dice (left) for Mouse (top) and Marmoset (bottom) datasets. Metrics computed per ontology level.

For the marmoset dataset, the same trends are observed (Fig. 6). Notably, labels and Nissl images transfer better to each other than to MRI. This is likely because these atlas labels were drawn over Nissl images and match their contrast and resolution, whereas MRI data has is significantly lower contrast and resolution. This indicates that appearance mismatch is a stronger predictor of performance than transformation direction. Cross-modality performance is also more symmetric than in mouse. At finer levels, performance again degrades substantially. This is consistent with increasing label sparsity and structural complexity.

### 3.2 Slice Localisation

We evaluate the impact of slice localisation on downstream registration by comparing predicted versus oracle slice selection on a held-out validation set of 256 samples per species. Localisation mean absolute error (MAE) is 0.95 ± 0.09 slices (47.5 ± 4.5 *µ*m) for mouse and 2.46 ± 0.13 slices (246 ± 13 *µ*m) for marmoset. The higher marmoset MAE reflects the smaller inter-slice anatomical variation of the marmoset atlas rather than under-training: performing twice as many epochs of training (essentially doubling training data) only marginally reduces this floor.

Despite this difference, end-to-end registration tracks oracle-slice performance closely. Across all modalities and both species, *Δ* Micro Dice (predicted − oracle) remains within − 0.04 at coarse and intermediate levels, and *Δ* HD95 within 2 voxels (Fig. 7). The level-7 dip persists under predicted-slice evaluation, confirming it reflects atlas structure rather than localisation error. Performance variability grows at the finest levels in marmoset due to the small number of structures.

**Fig. 7:**
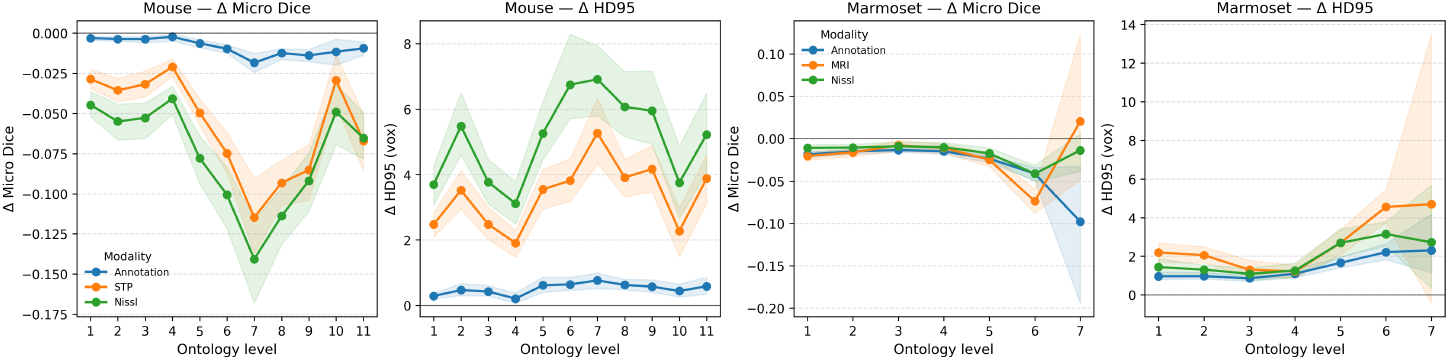
*Δ* Micro Dice and *Δ* HD95 (predicted minus oracle) across ontology levels for mouse (panels 1–2) and marmoset (panels 3–4).

### 3.3 Comparison with DeepSlice

We compared our pipeline against DeepSlice [6]as a baseline for mouse brain slice localization. Both methods were evaluated on an identical set of 200 synthetic coronal sections per modality (STP and Nissl) generated at 50 *µm* resolution in both affine-only and full (affine + non-linear) settings. To ensure a fair comparison, all metrics were computed in the original input space. Two levels of geometric distortion were tested by constructing two separate sets of synthetic deformations for our coronal slice: Affine (rotation/scale/shear) and Full (affine + non-linear deformation).

Our method consistently outperforms DeepSlice (Table 1), with the largest gains observed under non-linear deformation. When geometric distortion was produced with only affine transformations, improvements are moderate at coarse and intermediate levels, and diminish at fine levels for Nissl. The performance gap is particularly pronounced for STP, demonstrating strong cross-modality generalisation of our method.

**Table 1:**
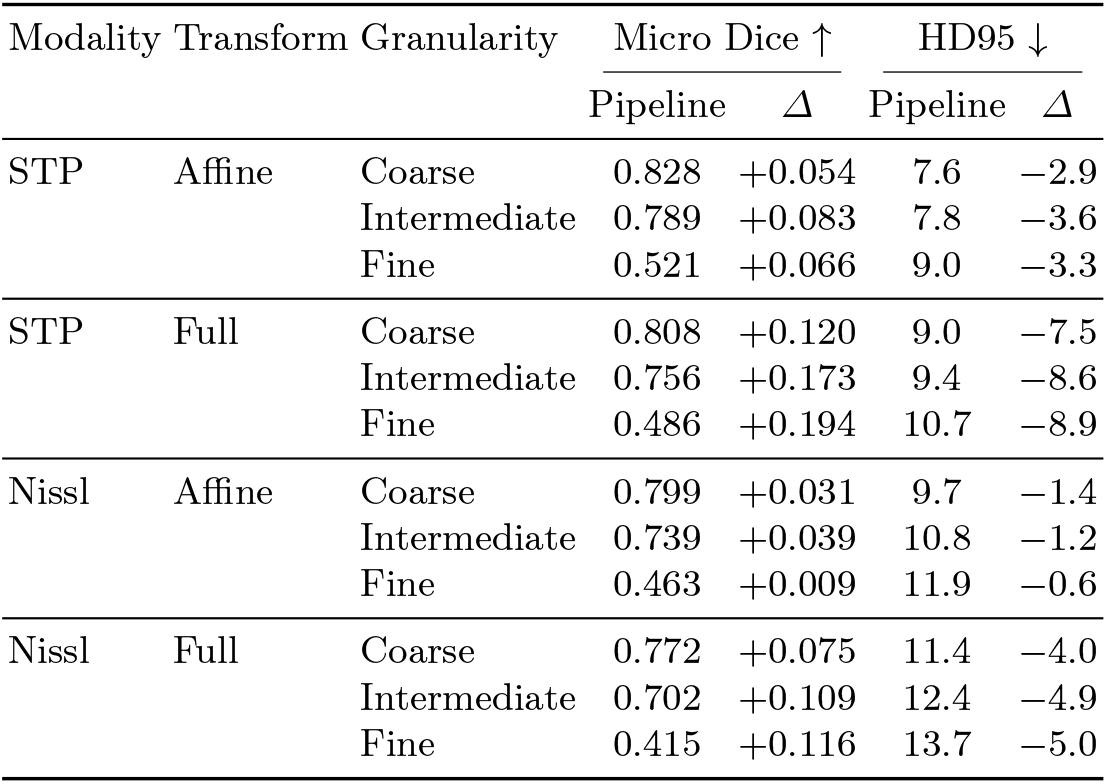
Pipeline vs. DeepSlice across ontology granularities. Positive *Δ* Micro Dice and negative *Δ* HD95 favour the pipeline. Results are averaged over paired bootstrap samples (*B* = 200).

Under full deformation, the advantage widens substantially across all granularities. Dice improvements increase with anatomical resolution. Corresponding reductions in HD95 confirm improved boundary alignment.

These results highlight that registration capability, rather than appearance matching alone, is critical under spatial distortion. DeepSlice, which lacks a deformable registration stage, degrades rapidly as deformation complexity increases. This validates that a learned deformation stage is essential for accurate cross-modality alignment, not just a small refinement.

## 4 Discussion

### 4.1 Summary

#### Registration quality and practical scope

Even though the pipeline must predict the atlas slice itself, it registers almost as accurately as when given the ground-truth slice (the *oracle*): at coarse and intermediate anatomical scales, Micro Dice stays above 0.75 (within 0.04 of the oracle) and HD95 within 3–4 voxels of oracle performance, across modalities and species. This corresponds to reliable delineation of major functional systems (e.g., thalamus, cortex, striatum), which is sufficient for most large-scale neuroanatomical analyses.

Performance degrades at fine scales, reflecting the intrinsic difficulty of aligning structures that span only a few voxels at 50 *µ*m resolution. This limitation likely arises from atlas resolution and boundary ambiguity rather than model capacity.

#### Robustness of registration to localisation error

Registration is robust to multi-slice localisation error. With localisation MAEs of 0.95 slices (mouse) and 2.46 slices (marmoset), end-to-end Dice remains within 0.04 of oracle performance at coarse and intermediate ontology levels across all modalities. The deformation field absorbs small residual offsets because adjacent atlas slices share similar anatomical structure, allowing locally smooth correction. This indicates that precise slice localisation is not the primary bottleneck for alignment accuracy in the regimes evaluated.

#### Cross-modality generalisation via progressive colour augmentation

The strong performance on STP and Nissl demonstrates genuine cross-modality generalisation. Progressive colour augmentation forces the network to rely on structural geometry rather than intensity cues, eliminating the need for explicit modality translation.

#### Versatility: atlas and pairwise alignment

Because each training pair is generated as two independent appearances of a common slice, the same network applies to two settings without retraining: aligning a single section to an atlas (the one-input pipeline evaluated here), and directly registering pairs of sections, for example adjacent serial sections, or differently stained sections of the same tissue. The model is agnostic to which image serves as the reference.

#### Fast, label-free registration

Registration runs in a single forward pass, reducing alignment time by orders of magnitude relative to iterative optimisation-based methods such as SyN [3] or LDDMM [5]. At the same time, the synthetic training strategy removes any need for manually labelled data, combining the speed of learned registration with the label-free generality of classical methods. This contrast-invariant, synthesis-based approach parallels SynthMorph [14], developed for 3-D human brain MRI. Here we show that the same principle extends to serial 2-D histological sections and generalises across organisms. Unlike SynthMorph, which performs image-to-atlas alignment only, our formulation additionally supports pairwise slice registration.

#### Accuracy across the full ontology

Rather than reporting accuracy for a small, fixed set of regions, we characterise it at every level of the brain ontology. This dense, hierarchy-wide evaluation is enabled by our synthetic training data, which supply exact ground-truth labels for every structure on every slice.

### 4.2 Limitations

#### Quantifying accuracy across ontologies

Reporting accuracy at every level of the ontology is itself (as opposed to choosing a fixed set of structures) a relatively novel way to quantify alignment accuracy, and the best methodology remains an open question. Important questions include “How to weight structures of very different sizes?”, “how to treat sparse or near-single-voxel regions?”, and “how to compare levels whose semantic granularity differs across species”. We regard our protocol as a step towards, rather than a final answer to, ontology-aware evaluation.

#### Coronal, in-plane assumption

The method assumes that input sections correspond to approximately coronal planar slices. We note that exactly the same method could be trained on sagittal or transverse slices. In practice, histological sections may be oblique due to acquisition variability. This could lead to systematic misalignment, as a 2-D registration model cannot correct out-of-plane tilt. While small tilts may not significantly affect coarse anatomical alignment, it may degrade accuracy in fine-scale or highly anisotropic regions. A natural extension is to estimate the section plane in 3-D and to model out-of-plane and in-plane rather than only in-plane deformation prior to registration. Our previous work designed classical iterative registration methods for this purpose [21,23], Deep-Slice (for example) uses an anchor-based formulation [6], and others reconstruct a full 3-D volume [16,20]. However some modern brain cell atlases, such as the Allen Institute’s spatial transcriptomics ABC atlas [27], consist of single section planes with larger gaps between them, such that matching to an appropriate section in standard orientation will be useful.

#### Resolution limits at fine scales

All experiments are conducted at 50 *µ*m atlas resolution. At this scale, fine anatomical structures occupy only a few voxels, making Dice and Hausdorff metrics highly sensitive to small spatial errors. This imposes a fundamental limit on achievable accuracy at the finest ontology levels. Higher-resolution atlases (e.g., 10–25 *µ*m) would provide a more stringent evaluation of fine-scale alignment.

#### The objective function

Our registration network was trained by directly predicting affine and velocity field transformation parameters by minimizing square error. As part of this approach we found that it was critical for velocity fields to be supported only on image gradients before smoothing (as described in [17]). As one would expect, our method was unable to predict displacements in homogeneous image regions where there are no visual cues. This nonidentifiability is related to the stabilizer of the image under the group action [22]. Other deep learning approaches have used alternative loss functions that may address the stabiliser directly. For example, SynthMorph [14] maximised Dice overlap between segmentations, and QuickSilver [26] minimised sum of square differences in image intensities. These two objects (segmentations and images) are both computed downstream of the transformation, and as such may come with additional sources of noise. However, tradeoffs between these sources of noise, and issues with identifiability are yet to be explored.

### 4.3 Cross-species generalisation and future directions

The mouse and marmoset experiments use identical network architectures and training procedures, differing only in the atlas volume and ontology used to generate training data. Comparable performance across both species suggests that the approach is genuinely transferable across species without species-specific engineering. Extension to other species (rat, macaque, human post-mortem) would require only an appropriate 3-D atlas and its annotation ontology, both of which are increasingly available through community atlas projects [8]. The addition of further modalities (e.g. NeuN staining, multiplexed fluorescence) is similarly straightforward within the existing colour augmentation framework. The synthetic appearance model could likewise be extended to emulate tissue-processing artifacts (e.g. tears, folds, missing tissue, and staining inhomogeneity) further narrowing the synthetic-to-real gap.

## 5 Acknowledgments

This work is supported by the National Institutes of Health grant number UM1 NS132173.

